# Development of a method for revertable CRISPR/Cas9-based mutagenesis in cell culture

**DOI:** 10.1101/2020.09.07.286187

**Authors:** Jonathon Walsh, Jonathan Eggenschwiler

## Abstract

CRISPR/Cas9 mutagenesis is a revolutionary tool for genetics in organismal and cell culture systems. One notable caveat with this system is the potential for phenotype-inducing off-target/background mutations. There has been considerable success in modifying the methodology to minimize these potential confounds. Here we have developed a tool to functionally demonstrate that a targeted mutation of interest is responsible for the phenotype observed. This approach creates revertable mutations in cell culture systems using CRISPR/Cas9-induced homology-directed repair (HDR) to insert a LoxP-flanked transcriptional stop sequence into an early intron of a target gene. This method has the potential to be used in multiplexed and inducible scenarios to restore gene function within a given experiment.

## Introduction

CRISPR/Cas9, a powerful tool for sequence directed mutagenesis, has revolutionized forward and reverse genetics. Breakthroughs in many genetic disorders have come from CRISPR/Cas9 genome editing in cell culture [1]. Recently, a human genetic disorder mutation was successfully corrected in viable human embryos [2]. With this technology, some limitations still exist. The incidence of off-target mutations is often the most discussed issue when utilizing CRISPR-based strategies and much effort is being put forth to overcome this issue [3]. The use of shorter gRNA sequences, Cas9 variants that have lower capacity for off target mutations, paired nickase Cas9, and catalytically inactive Cas9 fused to a cytosine deaminase to direct a single base changes have all been proven useful strategies for reducing the frequency of off target mutations [4–7]. All of these methods require whole-genome-sequencing strategies to detect the presence or absence of off target mutations. In addition to potential confounds resulting from CRISPR-induced off target lesions, the passage of cells in culture can lead to mutations through genetic drift and may be expanded within the culture when generating clonal lines that require extensive passaging [8]. Completely abolishing off-target effects of CRISPR/Cas9 mutagenesis is imperative for future use of this technology in treating human genetic disorders. For many applications, however, functionally benign off-target mutations may be acceptable as long as the phenotype under study can be shown to result from the desired locus-specific lesion introduced.

Several techniques have been used to circumvent issues with off-target mutations and *de novo* mutations generated by passaging cells, such as phenotype validation using multiple Crispr/Cas-induced genetic lesions in the same gene. This can be problematic since phenotypes may vary (expressivity/penetrance issues) among lines targeted with different guide RNAs due to variable loss of function. Alternatively, researchers have used complementation with wild-type expression constructs but this, too, may result in confounding results as overexpression constructs are often not under endogenous regulatory control and may cause phenotypes on their own.

Here we describe a method to functionally validate genotype/phenotype relationships in CRISPR/Cas9 mutagenesis experiments by confirming that phenotypes do not arise from off-target/background lesions. In addition, the modular potential of this method lends itself to multiplex mutagenesis and the ability to restore endogenous gene function within an experiment. This method utilizes CRISPR/Cas9 mutagenesis to insert, through homologous recombination, a LoxStopLox transcriptional stop cassette into an early intron of a gene to block transcription (**Fig. 1**). Since this insertion is in an intron, Cre reversion will restore transcription without disrupting the reading frame. The use of homology directed repair (HDR) to insert a relatively large DNA fragment into the genome makes simple and inexpensive strategies useful to probe the genome for off target insertions such as southern blotting. This is a unique system that allows Cre-induced complementation in the same cells that contain the genetic lesion, making this a very tightly controlled system for variation that may arise from differences in cell lineage using previously described methods.

**Figure 1:**
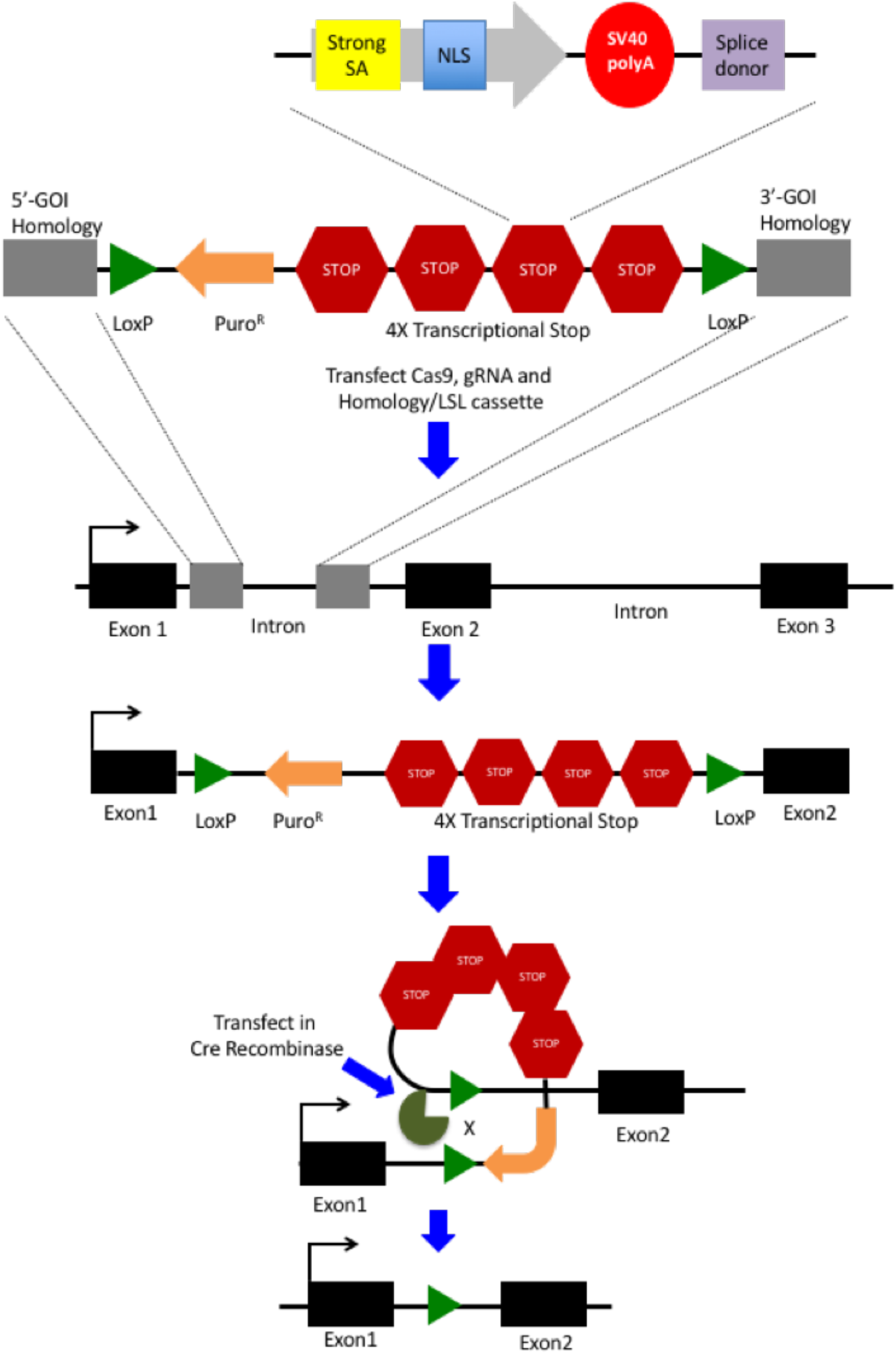
CRISPR-mediated insertion and Cre-mediated removal of the LoxStopLox transcriptional stop cassette. The LoxStopLox transcriptional stop cassette is flanked by LoxP sites for Cre mediated removal of the transcriptional stop sequences. A puromycin resistance gene driven by a PGK promoter is directly downstream of the 5’ LoxP site to be transcribed in the opposite direction of the target gene. Four tandem transcriptional stop sequences are positioned downstream of the resistance gene. Each stop sequence is comprised of an adenoviral splice acceptor sequence, a nuclear localization signal, a SV40 polyA termination sequence, and a splice donor sequence. This design ensures that transcription is effectively terminated, and if there is any leakiness, aberrant splicing or transport to the nucleus should ablate function of any translated protein. The homology flanked transcriptional stop cassette is inserted into the first intron of the gene of interest (*IFT172*) through HDR induced by a CRISPR-mediated double strand break. Successful HDR will stop transcription, creating a functionally null allele. Once a clonal cell line containing a homozygous insertion of the LoxStopLox cassette has been identified, the stop cassette is removed by transfecting a Cre expression vector. This will allow transcription of the gene of interest to proceed as normal.

As proof-of-principle, we have applied this approach to the mutagenesis of *IFT172* in mouse IMCD3 cells. IFT172 is a crucial component of the IFTB complex, which is required for intraflagellar transport, thus necessary for the formation of a cilium. Mutant mice and cell lines null for *IFT172* do not generate cilia [9,10]. This conspicuous phenotype makes it a suitable choice for assaying the utility of this method on an individual cell basis.

## Materials and Methods

### Guide RNA selection

The online tool ZiFiT was used to determine the guide RNA (gRNA) sequence [11,12]. A truncated 17-mer gRNA was used to minimize off target insertions [13]. Once selected, the guide sequence was obtained from IDT as a gBlock consisting of a U6 promoter, the gRNA sequence, and the gRNA scaffold [14].

### Generation of a homology sequence

60 base pairs of genomic sequence flanking the gRNA site was selected as the homology arms for homologous recombination. A homology construct was digitally generated using A Plasmid Editor (ApE) containing the 5-prime and 3-prime homology sequences, a restriction enzyme site flanking the homology sequences (Nru1), and two unique restriction enzyme sites separating the homology sequences (NotI/SacI). This entire cassette was also flanked by two more unique restriction enzyme sites (EcoRI/BamHI) (**Fig. 2**). The homology sequence was obtained as a gBlock from IDT and amplified with appropriate primers.

**Figure 2:**
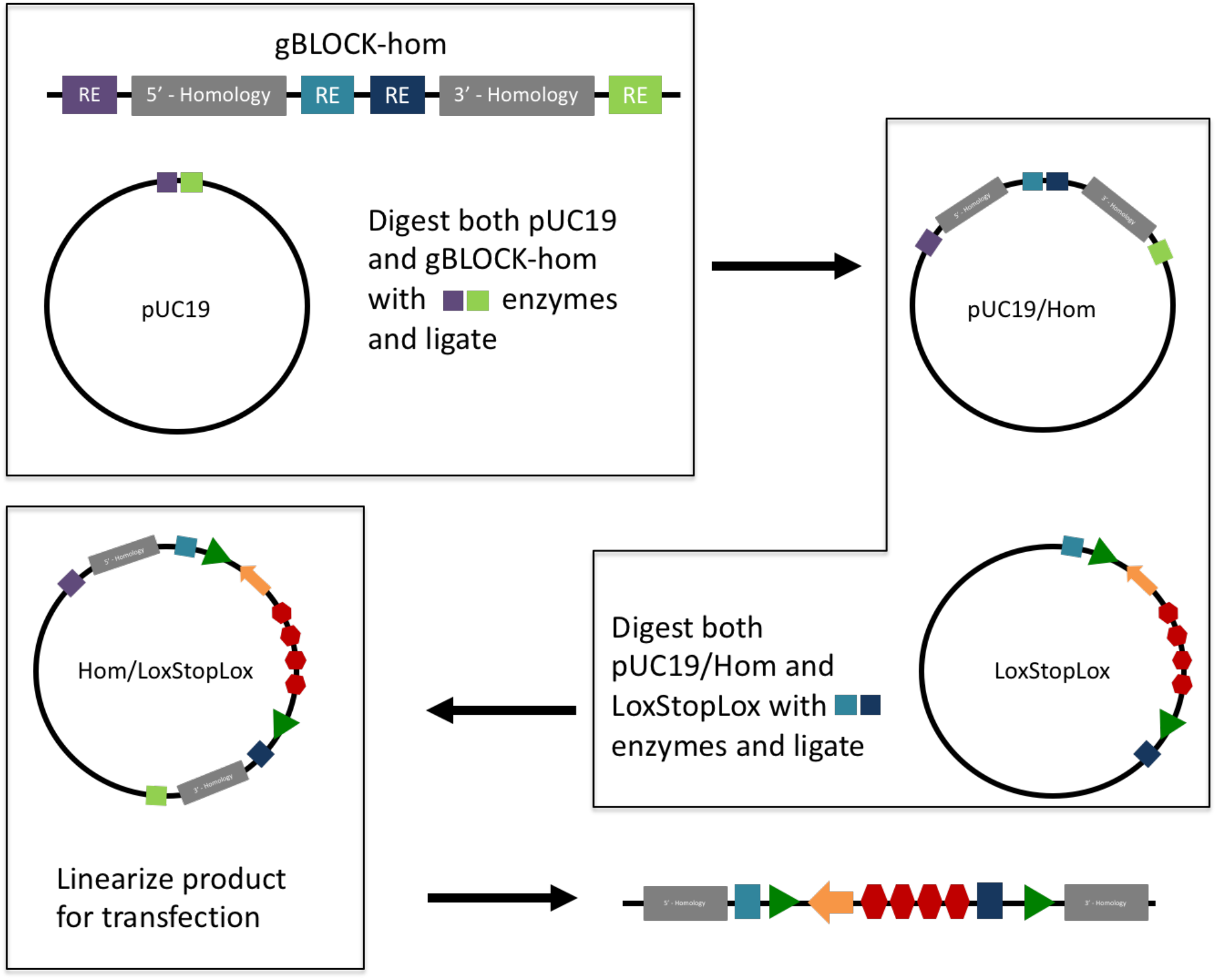
Cloning strategy for generating the homology-flanked LoxStopLox transcriptional stop cassette. A gBLOCK (IDT) was constructed containing the 5’ and 3’ homology sequences to induce HDR containing all the necessary restriction enzyme sites for cloning into a propagation vector (pUC19). The homology cassette and pUC19 were digested with the same restriction enzymes for directional ligation into the propagation vector. The pUC19/hom and LoxStopLox vectors were digested with the same two restriction enzymes (different from the initial step) to ligate the transcriptional stop cassette into the pUC19/hom vector. The pUC19/hom/LoxStopLox vector was digested with an enzyme to remove vector components.

### Cloning of the LoxStopLox-homology vector

The homology sequence was digested with EcoRI and BamHI and directionally cloned into the pUC19 cloning vector. The Lox-Stop-Lox TOPO vector was generated by Dr. Tyler Jacks and acquired from Addgene (plasmid # 11584). The LoxStopLox cassette was cloned into the pUC19/homology vector directionally using the NotI and SacI restriction enzyme sites (**Fig. 2**).

### Transfection and selection

Mouse IMCD3 cells were cultured in medium containing 10% FBS, 1X pen/strep, 1x GlutaGrow, and DMEM F12 (50:50). The cells were co-transfected in 60mm culture dishes at 60-80% confluence with PCR amplified gRNA, linearized LoxStopLox/homology cassette (NruI digested), and hCas9 plasmid (from Dr. George Church via Addgene, plasmid # 41815) using Lipofectamine 3000. The cells were cultured for 30 minutes in OptiMEM prior to adding the DNA/Lipofectamine complex and the medium was replenished with the usual growth medium after 8 hours. Cells were trypsinized, transferred to a 100mm culture dish, and placed under Puromycin selection (1ug/mL) 48-hours post transfection for 10 days.

### Generating clonal cell lines for PCR screening of insertions

After selection, cells were trypsinized and played at a limited dilution to obtain colonies derived from single cells. After 10 – 15 days of growth, 192 colonies were picked by incubating them in 20 mL of 5% trypsin/EDTA solution in PBS (5 mL of 0.25% Trypsin/EDTA in 95mL PBS) for 5 minutes at 37°C. colonies were gently aspirated using a micropipette and placed in a 96-well plate containing 150 uL of pre-warmed medium. Clones were cultured for 1 – 2 weeks. When clones were large enough, they were split into duplicate 96-well plates.

### Screening of clones

Three sets of primers were designed using NCBI Primer-BLAST: a set to amplify the wild-type sequence and two sets to amplify both the 5’and 3’ regions of the LoxStopLox homology cassette (**Fig. 3A**). DNA was extracted from one set of replicate 96-well plates. Medium was aspirated from cells and they were washed with PBS. 50 uL of 50mM NaOH was added to the cells and the plate was incubated at 95oC for 30 minutes. NaOH was neutralized with 10 uL of 1M Tris (pH 7.5). 1uL of this crude extract was used for PCR, performed with each of the primers to determine if the full length LoxStopLox cassette was inserted.

**Figure 3:**
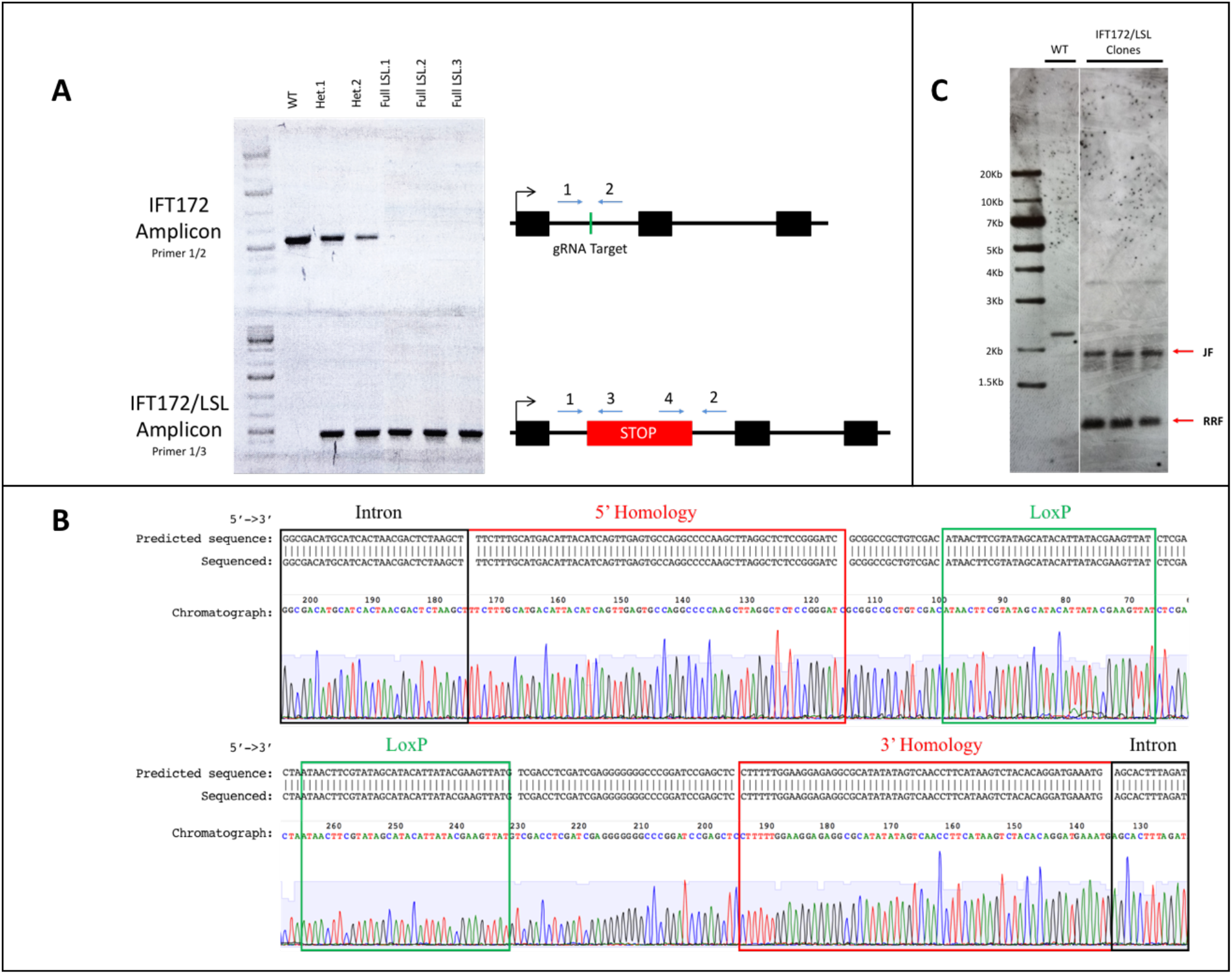
Genetic analysis of *IFT172/LoxStopLox* homozygous clones. **A)** PCR was used to identify clones with heterozygous and homozygous insertions of the full length LoxStopLox cassette. The WT control (primers 1 and 2) only amplified using primers flanking the endogenous site of mutagenesis and do not produce a product if the insertion is present. Genotyping of the insertion utilized one genomic primer and one primer homologous to part of the insertion sequence. Primers flanking both ends of the insertion (primers 1 and 3; 2 and 4) were used to ensure full length insertion. This genotyping design can differentiate between clones without an insertion, heterozygous for the insertion, and homozygous for the insertion. **B)** The PCR amplified fragment of the genomic hybrid sequence region at the 5’ and 3’ ends of the LoxStopLox homology cassette. PCR was amplified from genomic DNA from an *IFT172/LSL* clonal cell line. Sequence along the top is the predicted sequence based on manipulation of the sequence aligned to the actual sequence obtained. Boxes outline the aligned sequences and the chromatograms from the sequencing reaction. The black boxes surround sequence from the intron of *IFT172* upstream/downstream of the insertion. The red box surrounds the 60bp homology sequence that was used for HDR of the LSL cassette. Green box surrounds the LoxP site on the 5’ end of the insert for the purpose of Cre recombination. **C)** Genomic DNA of wild-type and *IFT172/LSL* clones were digested with AseI and probed with a DNA fragment that would identify any off-target insertions. Two bands were predicted based on the probe homology to the insertion. one band at 854bp demonstrates probe binding to three copies of the transcriptional stop sequence. The 1.9Kbp band represents the third copy of the transcriptional stop sequence that extends into the genomic sequence of *IFT172* from the 3’ end of the insertion. The 854bp band is ∼3 times more intense than the genomic band demonstrating no recombination events occurring within the repetitive sequence. A single band, not related to the insertion, was detected in the WT sample. The appearance of this band was variable, depending on the experiment (occasionally appearing in wild-type or mutant samples). A faint∼3.5kb band was observed in the homozygous clones. This corresponds in size to a predicted partial Ase1 digest fragment from the modified locus.

### Generating homozygous mutants from heterozygous clones

In a single round of mutagenesis, we identified heterozygous, but not homozygous, insertions. Homozygous mutants were generated from gene conversion by transfecting heterozygous clones with the hCas9 plasmid, the gRNA, and pTagRFP-C (Evrogen) as above. 48 hours post transfection, cells positive for RFP were isolated using a BioRad S3 flow cytometer. Clonal cell lines were generated and PCR screened as described above.

### Southern blot Analysis for off target Insertions

A DIG based DNA probe labelling kit was used to make a dsDNA probe for detection of the LoxStopLox insertion in the genome to verify the absence of secondary insertions. A gBLOCK was designed containing part of the transcriptional stop sequence which would not be present in wild-type genomic DNA. All 4 of the transcriptional stop repeats contain this sequence and are flanked by AseI restriction sites. The fourth repeat only contains a 5’ AseI restriction site with the downstream AseI restriction site extending into the genomic sequence. Thus, 2 bands are predicted in *IFT172/LSL* clones: a band at ∼1.9Kbp indicating a hybrid sequence containing one of the transcriptional stop repeats and *IFT172* genomic sequence and a band ∼854bp (3x more intense) representing the other three transcriptional stop repeats. Southern blotting was performed as instructed in the instruction manual for the DIG-High Prime DNA Labeling and Detection Starter Kit II (Roche Cat. 11585614910).

### Cre-mediated reversion of the LoxStopLox cassette

Upon verification that the cell lines homozygous for the LoxStopLox cassette were phenotypically null, the LoxStopLox cassette was removed by transfecting the cells with LV-Cre (from Inder Verma via Addgene, plasmid # 12106) using the previously described transfection protocol. For imaging purposes, cells were transfected on glass coverslips and fixed 48 hours after transfection in 4% PFA.

### qRT-PCR to verify loss of gene transcription

RNA was extracted from cells using the OMEGA E.Z.N.A. Total RNA Kit I. cDNA was synthesized using qScript cDNA SuperMix (QuantaBio). qRT-PCR was performed using PerfecTa SYBR Green FastMix (QuantaBio) and validated qPCR primers from IDT for *IFT172* mRNA expression and *β-actin* for internal control (PrimeTime).

### Immunofluorescence imaging of cells

Cells for imaging were grown on coverslips until 100% confluent and fixed in 4% PFA for 10 minutes. Cells were washed 3X-10 minutes in PBS. Cells were incubated in perm/block for 1 hour (10% heat inactivated goat serum, 0.1% Triton X-100, PBS).

Primary antibody dilutions were made in dilution buffer (1% heat inactivated goat serum, 0.1% Triton X-100, PBS) and cells were incubated in primary antibodies overnight at 4°C. Antibodies used were rabbit polyclonal against Arl13b (kindly provided by Dr. Tamara Caspary, Emory University) and a mouse monoclonal against acetylated α-tubulin (Sigma-Aldrich). Cells were stained with secondary antibodies along with DAPI diluted in dilution buffer. Cells were mounted on slides using Electron Microscopy Sciences FLUORO-GEL mounting medium. A Zeiss LSM 880 confocal microscope was used for imaging.

### Results

The homology/LoxStopLox cassette was created via a two-step cloning process (**Fig. 2**). Of 156 colonies analyzed, 13 (8.3%) showed partial insertion of the cassette and 3 (1.9%) had full-length heterozygous insertions. Homozygous clones were identified using a PCR genotyping strategy (**Fig. 3A**). Sequence analysis of both ends of the insertion confirmed seamless integration of the LoxStopLox cassette and confirmed the presence of functional LoxP sites (**Fig. 3B**).

qPCR analysis revealed little or no mRNA expression of the *IFT172* transcript (**Fig. 4**) in the homozygous clones. Southern Blot analysis demonstrated a single insertion of the LoxStopLox cassette (**Fig. 3C**). Cilia were completely ablated in three out of the five homozygous clones that were analyzed and transient transfection with a Cre recombinase expression plasmid restored ciliogenesis (**Fig. 5**). Two mutant homozygous clones (A4 and A5) showed rare (<1%) cells that generated cilia. This may have resulted from leaky read-through of the stop cassettes or contamination of clones with heterozygous cells.

**Figure 4:**
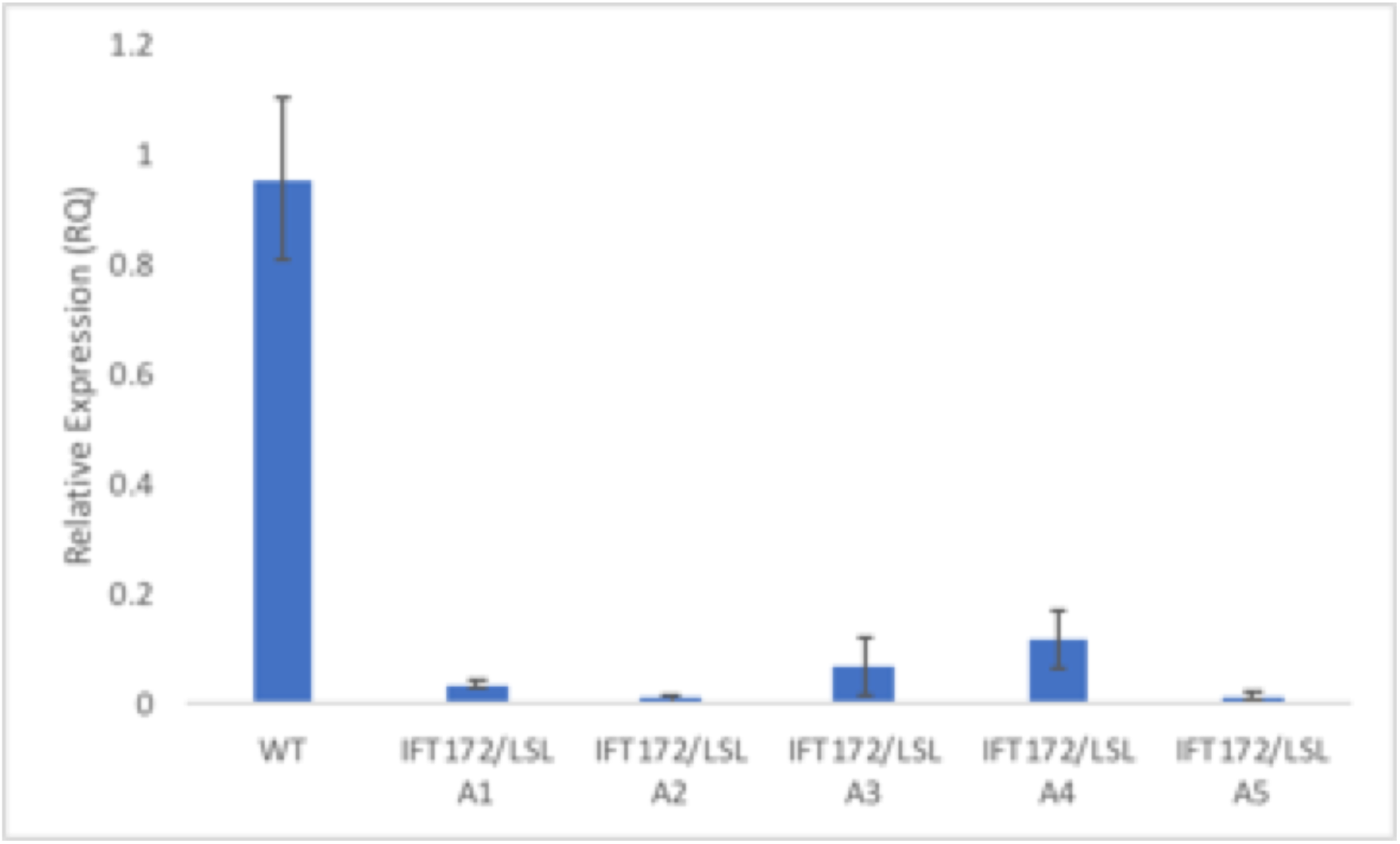
qPCR analysis demonstrates termination of *IFT172* transcription in *IFT172 LoxStopLox* homozygous clones. mRNA expression data from wild-type and five homozygous clones demonstrates a 90-99% reduction in *IFT172* expression. Expression of *IFT172* was normalized to that of *β-actin*.

**Figure 5:**
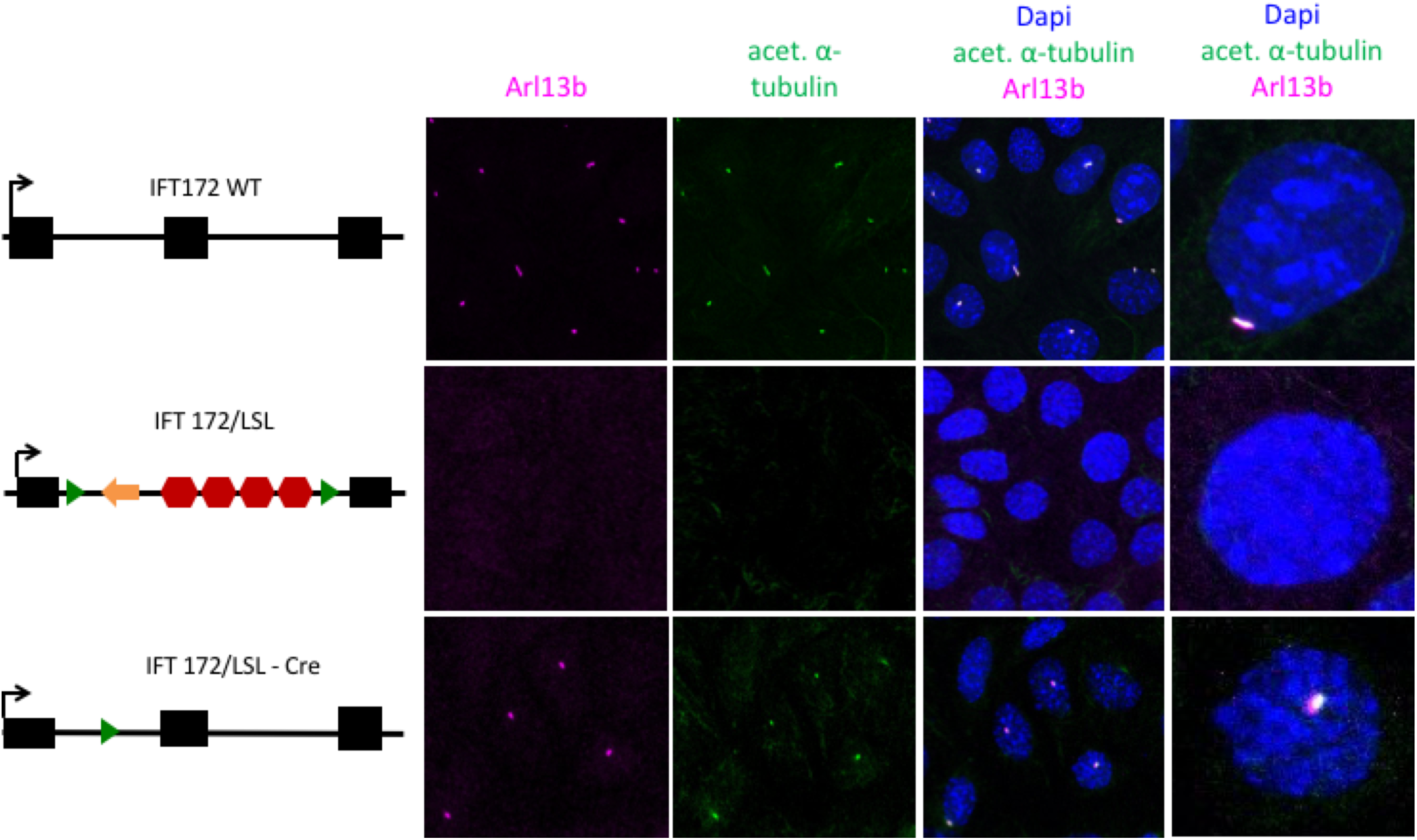
CRISPR-mediated LoxStopLox ablation of ciliogenesis and Cre-dependent recovery. Wild-type cells are induced to generate cilia by growing cells to 100% confluence. Cilia are co-labelled with acetylated α-tubulin (green) and Arl13b (Purple). Clones that are homozygous for the LoxStopLox cassette do not generate cilia. Upon transfection of these clones with a Cre expression vector, ciliogenesis was restored.

### Discussion

This method proves to be a reliable tool for generating revertable mutations in cell culture. This mutagenesis strategy was successful at disrupting *IFT172* expression/function, resulting in the failure to generate primary cilia. As the clones harbored only one insertion of the LoxStopLox construct (in the *IFT172* locus) and the block in ciliogenesis was reverted upon deletion of the cassette using Cre recombinase, the phenotype observed was clearly the result of the mutagenesis of *IFT172*. This was possible using a standard CRISPR/Cas9 system and very short (60bp) arms of homology on the HDR construct.

However, the frequency of full length insertions was roughly 2 in 100, which is representative of previously published results [15]. In the initial round of mutagenesis, only heterozygous mutations were identified. To generate homozygous mutations, a second round of mutagenesis of a heterozygous clone was required. For the second round of mutagenesis, the homology cassette was not included, and gene conversion was induced by the CRISPR/cas9-mediated double-strand break of the wild-type allele. Generation of homozygous mutants from heterozygous clones occurred at a frequency of roughly 20%.

Many innovations in CRISPR/Cas9 technology should prove useful for improving the efficiency of the mutagenesis. One study showed vastly improved HDR efficiency using a similar strategy but also removing all remnants of the vector backbone used to clone the homology sequences and increasing the length of homology to 800bp on each end [16]. In addition, synchronizing cells at M-phase using nocodozole has proven to increase HDR efficiency up to 33% in HEK293T cells [17] and small molecule inhibitors of NHEJ have similarly proven useful for increasing HDR efficiency in culture [18,19]. Implementing any of these strategies to our methods should increase efficiency of this method for generating revertable, functionally null mutations in cell culture.

In addition to combining this method with techniques for increasing HDR efficiency, this method allows for additional, novel experimental paradigms. One such use would be to make a CRISPR/Cas9-mediated LoxStopLox insertion in a gene in a cell line engineered for doxycycline or tamoxifen-inducible Cre activity [20,21]. With this paradigm, restoration of gene function in a temporal manner would be possible. In addition, with some basic modifications to the selective marker used on the LoxStopLox cassette, this method could be used to make double and triple mutants in the same cell line for studying genetic interactions at the cellular level. Combining this in a multiplexed format, with the ability to conditionally restore gene function would provide a sophisticated approach for addressing genetic interactions.

## Acknowledgements

The authors express their gratitude to two former undergraduate students, Ashley Dean and Kaley Desher, who helped with some of the experiments. The authors also wish to thank Dr. Julie Nelson (Director of the Cytometry Shared Resource Laboratory) and Dr. Muthugapatti Kandasamy (Director of the Biomedical Microscopy Core Facility) for technical assistance with flow cytometry and confocal microscopy.

